# Environmental exposure does not explain putative maladaptation in road-adjacent populations

**DOI:** 10.1101/095273

**Authors:** Steven P. Brady

## Abstract

While the ecological consequences of roads are well described, little is known of their role as agents of natural selection, which can shape adaptive and maladaptive responses in populations influenced by roads. This is despite a growing appreciation for the influence of evolution in human-altered environments. There, insights indicate that natural selection typically results in local adaptation. Thus populations influenced by road-induced selection should evolve fitness advantages in their local environment. Contrary to this expectation, wood frog tadpoles from roadside populations show evidence of a fitness disadvantage, consistent with local maladaptation. Specifically, in reciprocal transplants, roadside populations survive at lower rates compared to populations away from roads. A key question remaining is whether roadside environmental conditions experienced by early-stage embryos induce this outcome. This represents an important missing piece in evaluating the evolutionary nature of this maladaptation pattern. Here, I address this gap using a reciprocal transplant experiment designed to test the hypothesis that embryonic exposure to roadside pond water induces a survival disadvantage. Contrary to this hypothesis, my results show that reduced survival persists when embryonic exposure is controlled. This indicates that the survival disadvantage is parentally mediated, either genetically and/or through inherited environmental effects. This result suggests that roadside populations are either truly maladapted or potentially locally adapted at later life stages. I discuss these interpretations, noting that regardless of mechanism, patterns consistent with maladaptation have important implications for conservation. In light of the pervasiveness of roads, further resolution explaining maladaptive responses remains a critical challenge in conservation.

## Introduction

The global road network has expanded rapidly over the last half century (Canning 1998). Roads now cover some 64,000,000 km of the planet (Central Intelligence Agency 2013) and are projected to increase 60% by 2050 (Dulac 2013). Ecological consequences of roads are numerous and typically negative in effect. For example, roadkill causes an estimated one million vertebrate deaths per day in the United States. Habitat fragmentation spurs a suite of indirect effects (Forman and Alexander 1998) while runoff and leaching result in the deposition of a multitude of chemical contaminants into nearby habitats (Trombulak and Frissell 2000). Collectively, these effects extend well beyond the footprint of roads and are estimated to influence 19% of the land in the United States (Forman 2000).

Though the ecological effects of roads are well described, evolutionary outcomes remain poorly studied (Brady and Richardson in press). This is a critical gap in our understanding of road consequences because many of the negative effects of roads can be expected to act as agents of natural selection, causing populations to evolve. Specifically, natural selection occurs when variations of heritable traits (i.e. phenotypes) conferring relatively higher fitness are selected. That is, individuals expressing selected phenotypes survive and reproduce more successfully. The result of this evolutionary change is adaptation, comprising a shift in trait frequencies within a population toward phenotypes with higher fitness relative to the selecting environment. When selection pressures differ across local populations, divergent evolution can occur, resulting in local adaptation. Specifically, local adaptation is said to occur when populations evolve relative fitness advantages in their local environment compared to the fitness other populations experience in that environment (Kawecki and Ebert 2004).

Although any of the four mechanisms of evolutionary change (i.e. natural selection, genetic drift, gene flow, and mutation) can occur in roaded contexts, only natural selection results in adaptation, increasing the relative fit between populations and their environments. For example, evolutionary change by genetic drift can occur when roads sufficiently limit gene flow (Marsh et al. 2008). Much like natural selection, genetic drift differentiates populations. However, unlike natural selection, genetic drift is not expected to increase population fitness with respect to the environment. Thus, in the context of roads, natural selection is expected to increase the capacity of populations to tolerate negative road effects whereas other modes of evolution such as drift are not. Notably however, reduced gene flow can in some cases facilitate an adaptive response to selection by reducing the arrival of maladapted alleles (Garant et al. 2007; Richardson et al. 2016).

The small collection of studies that have investigated natural selection in the context of roads typically show that road-adjacent populations are adapted to road-specific selection pressures such as contaminants and road kill (reviewed by Brady and Richardson in press). This mirrors patterns of adaptation seen in many other contexts. For instance, reviews of reciprocal transplant studies indicate that local adaptation occurs in approximately 70% of cases (Hereford 2009; Leimu and Fischer 2008). In the context of conservation, the potential for local adaptation means that evolution can be a mitigating force contrasting the negative effects of environmental change.

Despite this relevance to conservation, local adaptation insights have traditionally been overlooked in applied ecological investigations (Hendry et al. 2010). Yet critically, local adaptation can occur quickly and across small spatial scales, matching both the pace and grain of environmental change and variation. Specifically, evolution can occur over handfuls of generations and across microgeographic distances (Hendry and Kinnison 1999; Richardson et al. 2014). This means that local populations can evolve divergently in traits and fitness across the landscape over both temporal and spatial scales that matter to conservation (Brown and Bomberger Brown 2013; Richardson and Urban 2013). Knowledge of this capacity for populations to evolve quickly and divergently has bolstered a recent and growing imperative to incorporate evolutionary perspectives in conservation (Carroll et al. 2014; Stockwell et al. 2003). Such work has further illuminated not only the pace and spatial scale of evolution, but also the potential for evolution to mitigate negative consequences of global environmental change (Hoffmann and Sgro 2011; Visser 2008).

### Describing local adaptation

Across populations, heterogeneous natural selection pressures coupled with limited gene flow can cause divergent evolution (Richardson and Urban 2013), such that populations become locally adapted to their local environments. Yet even in scenarios where adaptive traits are not heritable, phenotypic plasticity induced by the environment can generate a fitness advantage that is consistent with the pattern of local adaptation. Although these are distinct mechanisms by which adaptive outcomes can arise, they are not mutually exclusive. Moreover, plasticity itself can be heritable, while plastic and evolutionary changes can co-occur and interact. Further, plasticity may be relatively more widespread in the context of environmental change (Urban et al. 2014), and can serve as an important precursor to evolution, for example by inducing novel trait variation (Hua et al. 2015a). Therefore, from an evolutionary perspective, it is not particularly surprising that many populations facing environmental change show evidence for local adaptation (Hua et al. 2015b; Novak 2007).

Often, deciphering local adaptation is done experimentally within the context of reciprocal transplant experiments. Specifically, interacting fitness reaction norms—demonstrating a local population fitness advantage—is the diagnostic signature of local adaptation (Kawecki and Ebert 2004). Because this pattern of local fitness advantage can be generated either through evolutionary change (i.e. at the genetic level) or through phenotypic plasticity, multi-generation studies are typically required to discern the relative contribution of each of these mechanisms. This is because plasticity can be induced through parental environmental exposure independent of genetic variation (Rossiter 1996). Outside of model systems, parsing these mechanisms can be difficult. However, regardless of mechanism, local adaptation patterns highlight the scale and direction of fitness variation among populations responding to environmental change.

### Describing local maladaptation

Owing perhaps to a predominance of reported adaptive responses (Hereford 2009; Leimu and Fischer 2008), along with their potential to counter negative effects of environmental change (Gonzalez et al. 2013), local adaptation has become the default evolutionary hypothesis for populations responding to environmental change. Yet while local adaptation is indeed common, it is by no means the rule. That is, just as populations can become locally adapted by evolving a fitness advantage in response to changing environments, so too can they become locally maladapted, evolving a fitness disadvantage (e.g. Brady 2013; Falk et al. 2012; Rolshausen et al. 2015).

Whereas local adaptation is well described both conceptually and empirically (Kawecki and Ebert 2004; Savolainen et al. 2013), no formal framework exists for local maladaptation. Thus, compared to evidence for fitness advantages, evidence for fitness disadvantages is not only surprising, but also challenging to interpret (Crespi 2000; Hendry and Gonzalez 2008). Empirically, most examples of maladaptation are reported in the context of co-evolutionary dynamics (Thompson et al. 2001) and maladaptive gene flow (Lenormand 2002). More generally, maladaptation is described in terms of deviation from adaptive phenotypic peaks, wherein traits remain below some optimum fitness (Crespi 2000; Hendry and Gonzalez 2008). For example, a trait is considered maladaptive when other variants of that trait confer higher fitness in a given environment. This level of maladaptation can be assessed in the context of selection studies. However, maladaptation can also be thought of in terms of the fitness of a population. For instance, a population would be considered maladapted when its fitness is less than the fitness other populations achieve in that environment. Ultimately, natural selection operating on traits is the process that shapes maladaptive outcomes both in terms of the fitness of traits under selection and the population level response.

Here, I focus on the population aspect of local maladaptation in a manner analogous to that of local adaptation, while remaining complementary to the definition presented in terms of adaptive phenotypic peaks. Specifically, I define local maladaptation as occurring when populations evolve relative fitness disadvantages in their local environment compared to the fitness other populations experience in that environment. Mechanistically, maladaptation can result from several evolutionary processes. For example, when natural selection is strong and reduces population size, inbreeding depression can subsequently occur, resulting in a state of local maladaptation (Falk et al. 2012). Alternatively, high rates of gene flow from populations adapted to other environments can cause local maladaptation (Bolnick and Nosil 2007).

As with the pattern of local adaptation, evidence for local maladaptation can be caused by evolutionary and plastic change. For example in the context of a reciprocal transplant experiment, evidence of a local fitness disadvantage can be generated through genetic differences (true maladaptation) or induced in the form of plasticity. In the absence of knowledge regarding the relative contribution of these mechanisms, evidence of a fitness disadvantage can be referred to as ‘putative maladaptation’ (Crespi 2000). Regardless of mechanism however, local populations that respond maladaptively to environmental change have lower fitness than nearby populations, indicating an increased challenge to persistence.

Critically, there appears to be an emergence of studies reporting patterns of maladaptation in response to environmental change (Brady 2013; Christie et al. 2012; Rolshausen et al. 2015; Zimova et al. 2016). Despite the relevance of such outcomes to conservation, our knowledge about them remains limited and is challenged by the complexity of responses. For example, in the context of roads, evidence for local maladaptation is found in one species of amphibian despite evidence for local adaptation in another, which breeds and dwells in the very same habitats (Brady 2012; Brady 2013). That identical forms of environmental variation can drive divergent outcomes between related, cohabiting species suggests that local population level responses are complex and evolutionary responses to environmental change may be difficult to generalize.

Here, I focus on developing our understanding of a local maladaptation pattern that I previously reported in populations of the wood frog (*Rana sylvatica* = *Lithobates sylvaticus*) breeding in roadside ponds (Brady 2013). In that study, which I conducted in 2008, I used reciprocal transplant experiments across 10 populations to show that roadside wood frog populations (natal to ponds < 10m from a road) have 15% lower survival compared to road-naïve (hereafter ‘woodland’) populations. This survival disadvantage occurred both in the local roadside environment and in the transplant woodland environment, indicating that in addition to putative local maladaptation, roadside populations may be more generally depressed (i.e. so called “deme depression”). Similarly, experimental exposure to road salt—a widely applied road de-icing agent—caused increased mortality and malformations in roadside populations compared to woodland populations (Brady 2013).

The pattern of maladaptation in roadside wood frogs may be caused by a variety of mechanisms. These include 1) true maladaptation caused by evolutionary change 2) maladaptive environmental inheritance, and 3) negative carryover effects. Whereas true maladaptation dictates that heritable genetic variation is responsible for a reduction in fitness (Crespi 2000; Kirkpatrick and Barton 1997), maladaptive environmental inheritance (e.g. maternal effects) (reviwed by Rossiter 1996) and carryover effects are forms of phenotypic plasticity. In the case of maladaptive environmental inheritance, offspring phenotype would be negatively influenced by parental environmental exposure, independent of genetic effects. Carryover effects occur when an individual’s experience influences future outcomes (O'Connor et al. 2014).

From among these mechanisms, here I focus on experimentally testing for the presence of negative carryover effects. These effects may have mediated the maladaptation pattern previously reported (Brady 2013) because wood frog embryos used in that study were collected from natural breeding ponds within 36 hours of oviposition. During that time—over which embryos typically undergo 5 – 10 sets of cleavage—embryos were directly exposed to roadside water, which contains a suite of runoff contaminants (e.g. chloride ions from winter road salting) that reduce wood frog performance (Brady 2013; Karraker et al. 2008; Sanzo and Hecnar 2006). Conceivably, this period of early-stage embryo exposure to roadside water may have generated a negative carryover effect (Dananay et al. 2015; Hua and Pierce 2013) on survival, in a pattern matching local maladaptation. This would accord with other amphibian studies reporting that early exposure to osmotically stressful environments does not promote acclimation (Wu et al. 2014; but see Hsu et al. 2012). Instead, early exposure to osmotic stress can reduce performance at later stages (Hua and Pierce 2013), even after individuals are removed from the stressful environment (Wu et al. 2012). Relatedly, Karraker and Gibbs (2011) report that embryos of the spotted salamander (*Ambystoma maculatum*) experimentally exposed to road salt for nine days continue to lose body mass even after the exposure period ended. Hopkins et al. (2014) show that embryonic exposure to salt can compound larval mortality rate compared to larvae that were unexposed as embryos. In the context of resolving putative maladaptation reported in the previous study (Brady 2013), knowledge of a carryover effect would be viewed as experimental artifact, contrasting evidence of local maladaptation.

To evaluate this possibility, I used a highly replicated reciprocal transplant design conducted across 12 wood frog populations. To facilitate comparability, I reproduced the experimental time frame and context of the previous study (Brady 2013) but experimentally manipulated laying conditions of breeding wood frogs. This was done to test the hypothesis that early-stage embryonic exposure to roadside water influences survival. Specifically, I predicted that roadside embryos exposed to roadside water would show reduced survival relative to those embryos exposed to control water. This outcome would suggest that reduced survival rates are not maladaptive per se, but rather comprise a direct environmental effect that is induced by exposure during the early embryonic period (i.e. a carryover effect). To control for potential parental effects this system, I also evaluated the influence of adult body condition on offspring survival. Further, I examined family level variation in survival to gain a preliminary understanding of whether larval performance might be heritable. Finally, to provide context for interpreting experimental results, I conducted field surveys to estimate population size and quantify a suite of environmental variables that influence amphibian performance and distribution.

## Materials and Methods

### Natural history

The wood frog is widely distributed throughout eastern North America and much of Canada, with a range extending from within the Arctic Circle to the southeastern United States. Breeding is explosive, and within the study region populations reproduce in late March/April, when adults migrate from upland terrestrial habitat to breed in ephemeral breeding ponds. Across this study site, populations typically breed synchronously within several days of one another, when each female oviposits one egg mass containing approximately 800 eggs. Embryos develop over two-three weeks before hatching, and continue to develop as aquatic larvae throughout spring and early summer until they metamorphose into terrestrial juveniles.

### Site selection

I selected populations from each of six roadside and six woodland ponds (hereafter referred to as ‘roadside deme’ and ‘woodland deme’). Ponds are located in the northeastern U.S. (Fig. 1 inset) within and adjacent to the Yale Myers Forest, characterized by large swaths of native trees and low human population density. All ponds studied in 2008 (Brady 2013) were included in the present study. Two additional ponds from a related study of spotted salamander responses to roads (Brady 2012) were also included for increased sample size. Detailed selection methods are described elsewhere (see Brady 2012, 2013). Briefly, roadside ponds contained the highest conductivity (μS) (an indicator of road salt runoff) among breeding ponds in the region (Fig. 1 inset). Each roadside pond was paired with a woodland pond located > 200 m from the nearest paved road, and characterized by complementary abiotic conditions to minimize confounding variation. Across all ponds, inter-pair distance ranged from 880 – to 6060 m apart.

**Figure 1:**
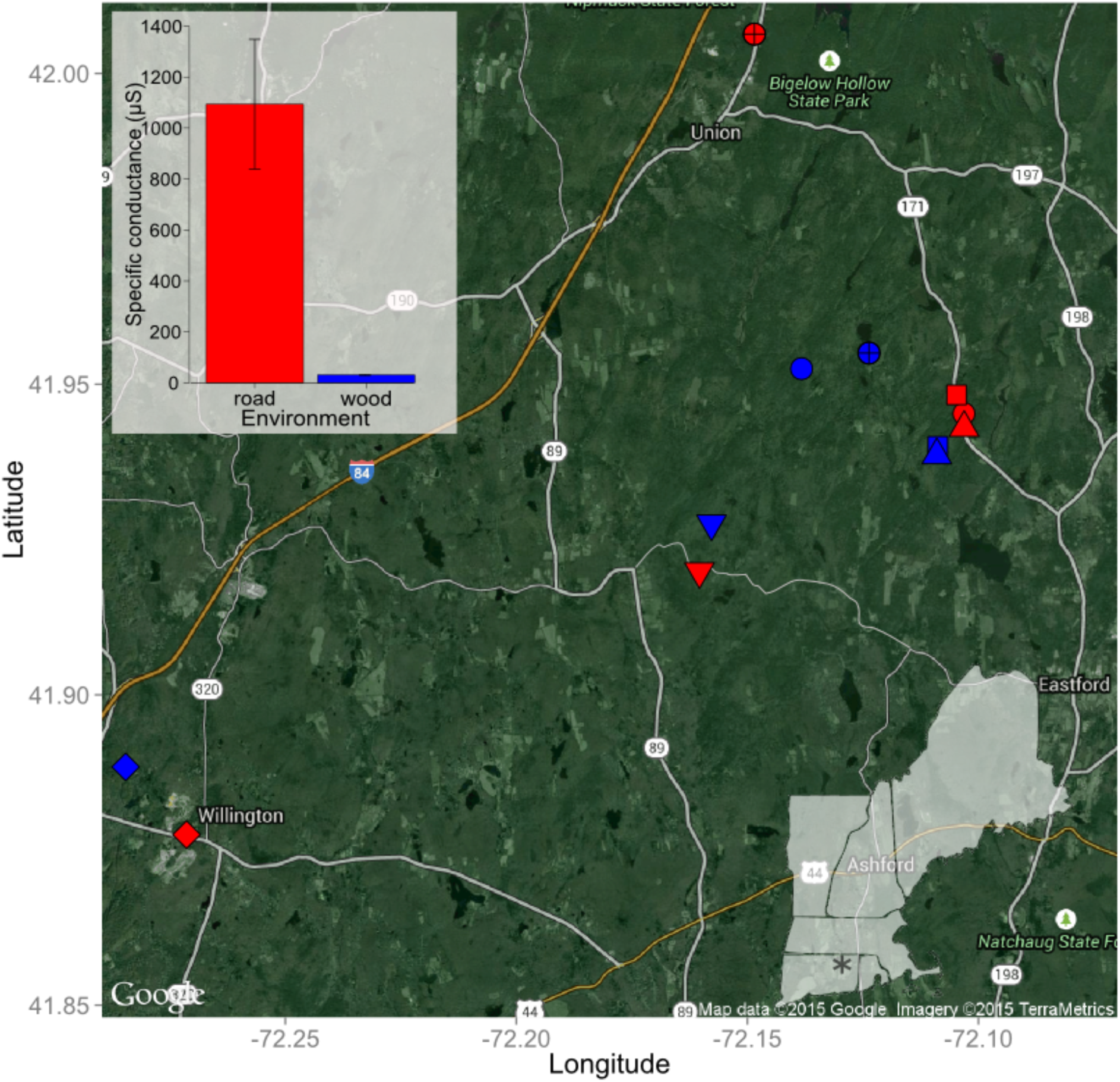
Study region and reciprocal transplant design. Location of 12 ponds comprising reciprocal transplant are shown on a map of the region. Red symbols indicate roadside ponds; blue symbols indicate woodland ponds. Each roadside-woodland pond pair comprising a deme level transplant shares a common symbol shape. Interstate highway (I-84) and on/off-ramp infrastructure is indicated in yellow. Primary roads are heavily shaded, while secondary roads are lightly shaded. Bar graph inset shows mean specific conductance (µS) (± 1 SE) for roadside (red) and woodland (blue) ponds. For roadside ponds, conductivity is shown as the average of surface and bottom values. This is because specific conductance in the bottom waters of roadside ponds is nearly twice that of surface water on average. No such vertical gradient exists in woodland ponds.

### Reciprocal transplant background and experimental design

For the present study, I used a reciprocal transplant experiment conducted in spring 2011 to evaluate survival, growth, and development of aquatic stage wood frogs. The key difference in the design of this experiment compared to the one previously reported (Brady 2013) is that here the natal environment was controlled for two days. In the previous experiment (conducted in spring 2008), embryos were collected out of ponds from naturally laid egg masses within 36 hours of oviposition, and were thus exposed to natal pond water for up to 36 hours. In the present study, I captured adults on their inbound breeding migration and controlled breeding so as to manipulate the natal environment. Thus, this experiment was composed as a 2 × 2 × 2 factorial design to test the interacting effects of deme, environment, and embryonic exposure on survival, development, and size. This design is analogous to the established ‘genotype by environment’ (i.e. G × E) framework used to test for local adaptation (Kawecki and Ebert 2004), with the additional term of embryonic exposure included to evaluate whether pond water influences the G × E outcome. In the G × E framework, interaction effects on fitness reflect differential responses among genotypes exposed to a common environment, indicative of population differentiation. The nature of this interaction provides inference into adaptation (see Kawecki and Ebert 2004) or maladaptation. For example, local adaptation is indicated when the local population—as compared to the foreign population—shows evidence of higher fitness within the local environment (Kawecki and Ebert 2004).

I used partially encompassing drift fences to collect adult wood frogs on their inbound breeding migration to each pond. Captured adults were measured for snout-vent length (SVL) and mass, and then paired to breed in 5.1 L plastic containers (33 × 20 × 11 cm), filled with 1 L of either respective pond water or spring water and placed on a slope at the edge of the pond. Breeding enclosures were monitored daily. Following oviposition, adults were released while embryos were maintained in their container for a two-day incubation period, mimicking direct natal environmental exposure reported previously (Brady 2013) and comprising the embryonic exposure treatment in this study (i.e. pond water versus spring water). Following incubation, I separated from each egg mass two clusters of ca. 80 embryos. Each of these clusters was stocked into its own separate enclosure (36 × 28 × 16 cm) within both the natal and complementary transplant ponds [Figs. 1 and Online Resource 1; enclosure details provided in Brady (2012)]. Thus, within a pond, each enclosure was stocked with full sib individuals from one unique family. Enclosures were subdivided such that each enclosure housed one local and one transplant family. Each enclosure comprised a block, while each subdivision comprised the experimental unit.

In total across 12 ponds, I stocked 327 experimental units (each with ca. 80 individuals), with a median of 14.5 unique families represented per population and a median of 8 unique families per population per treatment. Across all breeding containers, the date of oviposition ranged from 04 – 17 April and did not differ by deme (*P* = 0.169). I used a dissecting stereoscopic microscope to estimate the development stage of each egg mass upon stocking. At the conclusion of the experiment, when all eggs had either hatched and reached feeding stage (hereafter ‘hatchling’) or died, I estimated survival for each experimental unit. I haphazardly selected ca. half of these units (n = 176) to estimate SVL and development stage (Gosner 1960). I targeted 20 tadpoles per unit (actual mean = 18.0 ± 0.38 SE), staging and measuring a total of 3168 tadpoles across all experimental units.

### Characterizing population size and the environment

I waded through each pond after the completion of the breeding event to visually survey the number of wood frog egg masses as an estimate of population size. I also measured seven environmental characteristics associated with amphibian distribution and performance. Specific conductance, dissolved oxygen, pH, and wetland depth were measured once during the experiment, while temperature was measured every thirty minutes using deployed temperature loggers. Because of a vertical halocline present in roadside (but not woodland) ponds (Brady 2012), I measured specific conductance at both the top and bottom of the water column in roadside ponds. In 2008, global site factor—a measure of solar radiation reaching the pond—was calculated from hemispherical photographs, while wetland area was estimated from visual rangefinder measurements. Chloride was measured at eight ponds (four roadside, four woodland) using liquid chromatography (Brady 2012).

### Statistical analyses

All analyses were conducted in R V. 2.15.0 (R Development Core Team 2012). I used the package *lme4* to compose a suite of mixed effects models to estimate offspring performance variables (i.e. survival, development stage, and SVL) across the interaction of genotype × environment × embryonic exposure. Initial models differed in random effects structure; standard AIC selection criteria were applied to select the most parsimonious model. Survival was analyzed as a bivariate response of successes and failures using a binomial family with a logit link. Candidate random effects included pond pair, experimental block, family, and experimental unit (included to account for overdispersion). Inference for survival was conducted with MCMC sampling using the package *MCMCglmm*. The models for survival were composed with and without a covariate for adult body condition, which was estimated as the quotient of adult mass divided by adult SVL. Development stage and SVL were each analyzed as a univariate normal response and inference was based on Satterthwaite approximated degrees of freedom. Further, I examined whether survival, SVL, and development stage varied at the family level. This was done using a chi-square comparison of the model selected for inference with a model also containing a random effect term for family. I used MANOVA to evaluate the suite of abiotic variables characterizing the environment. Responses measured more than once at each pond were averaged. I used a linear model to evaluate the influence of pond type (roadside vs. woodland) on population size. Specifically, egg mass abundance was divided by pond area (i.e. egg mass density) because abundance varies with pond size (Karraker et al. 2008). Egg mass density was then log-transformed to meet model assumptions. All data for this study are available at: to be completed after acceptance.

## Results

### Reciprocal transplant: survival

Early exposure had no effect on survival (Posterior mean = 0.070, HPD_95%_ = -0.320 to 0.496, *P_mcmc_* = 0.756), nor did it interact with G × E to influence survival (Posterior mean = 0.418, HPD_95%_ = -1.190 to 2.190, *P_mcmc_* = 0.641). Survival varied across the G × E interaction (Fig. 2; Posterior mean = -1.349, HPD_95%_ = -2.092 to -0.549, *P_mcmc_* = 0.004). Within the roadside environment, survival was 29% lower for the roadside deme compared to the woodland deme (Posterior mean = -1.072, HPD_95%_ = -1.617 to -0.555, *P_mcmc_* < 0.001). Specifically 46% of roadside embryos compared to 65% of woodland embryos survived in the roadside environment. Survival was highest in the woodland environment (70%) and did not differ between demes (Posterior mean = -0.038, HPD_95%_ = -0.454 to 0.451; *P_mcmc_* = 0.889). Relative to the woodland environment, survival in the roadside environment was reduced by 7% for the woodland deme (Posterior mean = -0.706, HPD_95%_ = -1.254 to -0.170; *P_mcmc_* = 0.015) and 34% for the roadside deme (Posterior mean = -1.820, HPD_95%_ = -2.269 to -1.293; *P_mcmc_* < 0.001). There was no effect of female body condition on offspring survival (Posterior mean = -0.8609, HPD_95%_ = -5.5648 to 3.6437; *P_mcmc_* = 0.704), nor did inclusion of this term change inference into survival across the G × E interaction (P = 0.011). Similarly, although male body condition affected offspring survival (Posterior mean = 7.722, HPD95% = 0.427 to 14.040; *P_mcmc_* < 0.026), it did not qualitatively influence the effect of G × E on survival (P = 0.004). Finally, survival varied with respect to family (*Chi-square* _9,1_ = 4.246, P = 0.039).

**Figure 2:**
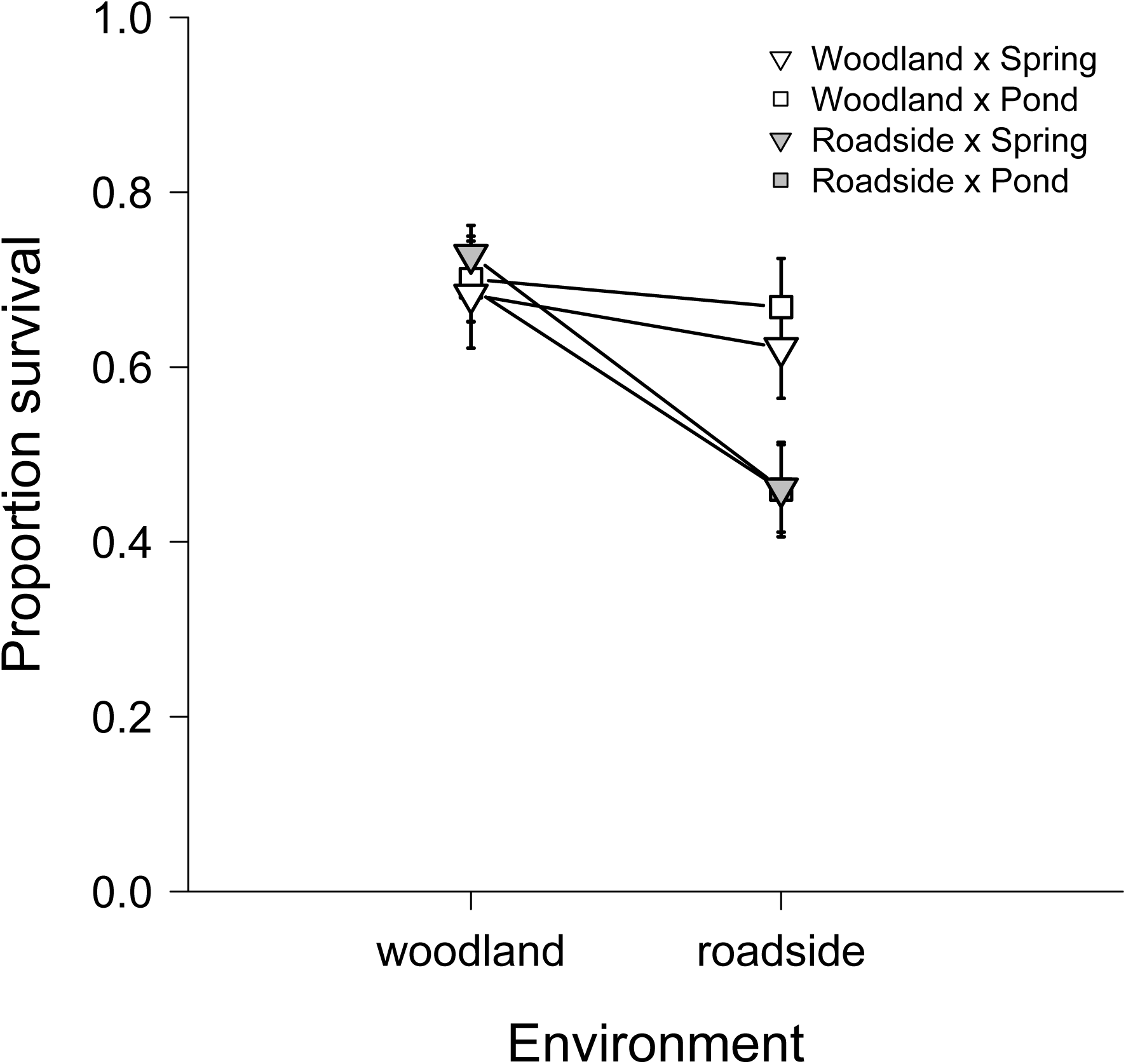
Survival across the G × E interaction and with respect to embryonic exposure. Proportion survival is shown as the mean from each experimental unit averaged (± 1 SE) across all experimental units (N = 327). Each experimental unit was stocked with embryos from a single family. Across the reciprocal transplant, each unique family of embryos was stocked into two experimental units: one placed in the local site and one placed in the transplant site. Open symbols represent the woodland deme while shaded represent the roadside deme; triangles indicate spring water treatment while squares indicate pond water treatment. Thus, ‘Woodland × Spring’ indicates woodland deme exposed to spring water treatment whereas ‘Woodland × Pond’ indicates woodland deme exposed to pond water treatment.

### Reciprocal transplant: hatchling development stage and size

Final development stage varied across the G × E interaction (*F* _1,68.38_ = 10.500, *P* = 0.022), and marginally with respect to the main effect of treatment (*F* _1,121.96_ = 2.958, *P* = 0.088). Among possible contrasts, development stage differed only for the woodland deme within the roadside environment and with respect to embryonic exposure (Fig. 3, panel a). Specifically, final development stage was 3.3% greater for woodland animals exposed to spring water versus pond water, and reared in the roadside environment (*F* _1,23.2_ = 5.078, *P* = 0.034). With regard to hatchling size, there was marginal evidence for a three-way interaction effect of G × E × embryonic exposure (*F* _1, 128.27_ = 3.327, *P* = 0.071). Hatchling size only differed for the woodland deme within the roadside environment and with respect to embryonic exposure (Fig. 3, panel c). Specifically, hatchlings from the woodland deme that were reared in the roadside environment were 10.7% longer when exposed as embryos to spring water as compared to pond water (*F* _1,21.90_ = 4.02, *P* = 0.057). Both SVL (*Chi-square* _7_,_1_ = 2248.0, P < 0.001) and development stage (*Chi-square* _8,1_ = 346.12, P < 0.001) varied at the family level.

**Figure 3:**
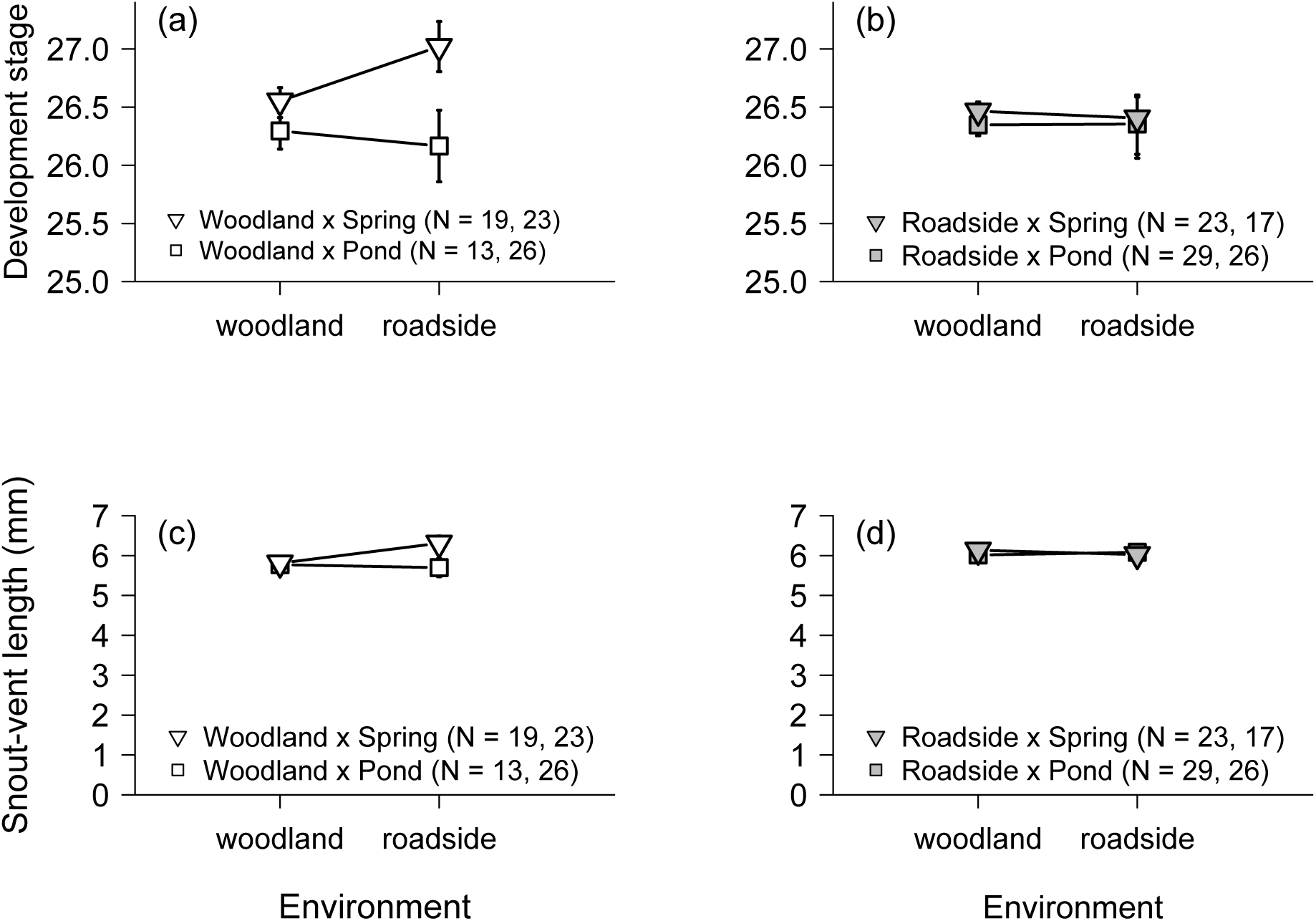
Development stage and snout-vent length (SVL) across the G × E interaction and with respect to embryonic exposure. Development (panels a and b) comprises mean Gosner (1960) stage. SVL (panels c and d) is a measure of larval body length. All responses are shown as the average (± 1 SE) of the means of each experimental unit. That is, measurements from all sampled larvae within each experimental unit were first averaged prior to then taking the overall mean across all experimental units for each treatment. For each treatment, two sample sizes (one for the roadside environment, one for the woodland environment) are shown in the key. Symbol descriptions are identical to those provided in Fig. 2.

### Population size and the environment

Egg mass density ranged from 0.019 to 0.234 per square meter and did not differ by deme (*P* = 0.712; Online Resource 2). The multivariate response of environmental variables differed across environment type (*Posterior mean* = 1.324, *P_MCMC_* < 0.001). Among these, follow-up univariate mixed models indicated that only specific conductance differed with respect to environment type (*F* _1,10.11_ = 31.99, *P* < 0.001 [Fig. 1 inset]; all other *P* > 0.458). Additionally, there was a vertical gradient in specific conductance in roadside ponds: values at the bottom of the ponds (where larvae frequent) were nearly twice that of the top of ponds (where embryos are typically laid). Specifically, in roadside ponds, specific conductance averaged 1428 μS (95% CI: 390.0, 2466.5) at the bottom of the water column and 758 μS (95% CI: 429.9, 1086.5) at the top of the water column, as compared to 31 μS (95% CI: 26.0, 36.3) in woodland ponds. Thus compared to woodland ponds, on average specific conductance of roadside ponds was 46 times higher at the bottom and 24 times higher at the top.

## Discussion

Consistent with previous findings (Brady 2013), roadside populations grown in their natal ponds survived at lower rates compared to populations transplanted there from nearby woodland ponds (Fig. 2). This pattern accords with local maladaptation. Moreover, there was no effect of the experimental environment experienced during the two days following oviposition. Thus, regardless of whether roadside embryos were conditioned in spring water or natal pond water, they experienced an equivalent survival disadvantage in their local environment compared to embryos transplanted there from woodland populations. I therefore found no support for the hypothesis that early embryonic exposure causes a carryover effect on survival in a manner that could explain the previously described maladaptation pattern (Brady 2013). Further, the survival disadvantage of the roadside deme in roadside ponds was not qualitatively influenced by variation in adult body condition. This adds confidence to the conclusion that the survival disadvantage depends on the G × E interaction, and adds support to the possibility that roadside populations are locally maladapted. More broadly, that the pattern of local maladaptation in this study system is now reported across multiple years and populations suggests that this phenomenon may be a generalized consequence for wood frogs breeding near roads.

Ultimately, the fitness consequences of this survival disadvantage depend on whether this effect persists into later life history stages. For example, a variety of processes such as density dependence in juveniles might mediate this survival pattern across life history stages, potentially offsetting the disadvantage. Unfortunately, the relationship between larval survival and population fitness in the wood frog is not well described, and is likely to be complex, varying across contexts such as density dependence and environmental conditions (Berven 2009; Dananay et al. 2015). I therefore discuss this survival disadvantage under different assumptions concerning the relationship between larval survival and population fitness. I first assume that the survival disadvantage in the roadside deme bears a negative effect on relative fitness; I then discuss the implications of this pattern when this assumption is relaxed.

That the survival disadvantage shown here occurred for embryos collected from a controlled breeding environment (i.e. spring water) suggests that this effect is parentally mediated. Assuming that survival is positively correlated with fitness, several potential mechanisms could explain how adult wood frogs mediate this maladaptive survival pattern on offspring. First, these results remain consistent (though are not conclusive) with true local maladaptation. This would imply that the roadside deme is genetically differentiated from the woodland deme, and that these differences are linked to a fitness disadvantage. This possibility is supported by family level variation characterizing SVL, development stage, and survival, indicating that these traits may be heritable, and can evolve in response to selection (Falconer and Mackay 1996). Local maladaptation could arise in several ways. For instance, maladaptation can result through the process of intense selection (e.g. for contaminant tolerance) decreasing populations to sizes small enough to cause inbreeding depression or drift (Falk et al. 2012). Likewise, maladaptation could potentially arise through novel maladaptive genetic variation originating by exposure to roadside contaminants acting as mutagens (Tchounwou et al. 2012). Intriguingly, this possibility raises the corollary that even if selection acted against such maladaptive genetic variation, for example favoring migrant wood frogs, the persistent nature of both past and ongoing contaminants in the roadside environment might result in a steady supply of maladaptive alleles in the population via mutagenic effects on each new generation.

Second, these results are consistent with the analogous phenomenon of maladaptive environmental inheritance. Unlike maladaptation, maladaptive environmental inheritance requires that a fitness disadvantage is caused by inherited environmental (not genetic) effects (Rossiter 1996). If this were the case, the survival disadvantage in offspring would be the result of parental environmental exposure, as is often described in terms of maternal effects. For example, this might occur as a result of increased parental stress (Saino et al. 2002; Tennessen et al. 2014). Alternatively, exposure to runoff contaminants associated with the roadside environment might induce a survival disadvantage (Metts et al. 2012; Todd et al. 2011). Distinguishing between these two mechanisms (i.e. maladaptation and maladaptive environmental inheritance) requires knowledge of whether genetic differentiation has occurred between woodland and roadside demes. Ultimately, such insights can be gained through multi-generational studies designed to control the parental environment and subsequent parental effects.

It is also useful to consider these results through the lenses of plasticity and evolutionary constraints, asking what might be limiting the roadside deme from an adaptive response. For example, given the relatively high survival capacity demonstrated by the woodland deme in the roadside environment, we might expect plasticity to enable a similar survival rate of the roadside deme. Yet although ubiquitous, plasticity is not limitless, and carries with it costs (Van Buskirk and Steiner 2009). Alternatively, adaptation in roadside populations may be constrained by presumably variable and strong selection regimes. In addition to the inbreeding effects mentioned above, such dynamics can result in maladaptation through lag effects (Crespi 2000). Indeed, variable and intense selection regimes have recently been suggested as potentially more important than costs of plasticity as factors limiting adaptation (Murren et al. 2015). Finally, potential adaptation to terrestrial pressures (such as road kill) might constrain adaptation to the aquatic environment.

Regardless of which mechanisms might be responsible for the survival disadvantage, the presence of either maladaptation or environmentally inherited maladaptive performance would hold similar and important inference for conservation. First, the negative survival effect of roadside ponds is more severe for the populations living there than for populations located away from the road. Thus, the populations that are most susceptible to road effects are also the least tolerant. Moreover, the cost of roadside dwelling is substantial, as evidenced by the woodland deme’s relatively high survival rate in the roadside environment. Another point to consider for conservation concerns the persistence of this pattern and its implications for restoration. Notably, in the previous study (Brady 2013), the survival disadvantage of the roadside deme occurred not only in natal ponds, but also in woodland ponds. Coupled with the contrasting outcome here—in which woodland pond survival is equivalent between demes—these results suggest that responses may be temporally variable. Yet even in years such as the one reported here where the roadside deme performs as well as the woodland deme in the woodland environment, the pattern of local maladaptation to the roadside environment persists.

Taken together, these findings suggest that a reversal of the survival disadvantage may be difficult even if management interventions are made to ameliorate the severity of the roadside environment. For example, in some years, survival following restoration efforts might improve while in others the survival disadvantage might persist. Moreover, if the survival disadvantage is caused by true maladaptation, genetic change may entirely preclude the reversal of negative effects. Further, without management intervention, reversal of negative survival might in some cases require a decoupling of the contemporary deme from future generations. Specifically, in line with the potential mechanisms (i.e. genetic versus environmental inheritance), this decoupling would require either genetic based adaptation (and thus population differentiation), or persistent migration from woodland populations bearing no previous exposure to the roadside environment.

Because my results do not provide inference into later life history stages, they should be interpreted cautiously as evidence for maladaptation. As mentioned above, ecological and biological contexts such as density dependence (e.g. Vonesh and De la Cruz 2002) and tradeoffs across life history stages (Dananay et al. 2015) can interact with and mediate the relationship between larval survival and fitness. Thus, if we relax the assumption that the negative relative survival of the roadside deme is correlated with fitness, a suite of alternative mechanisms can explain these patterns. For example, decreased survival at early larval stages might be offset by increased survival later in life, or by increased investment in fecundity, such that on the whole, roadside populations are actually locally adapted to road adjacency. While later stages of larval development do not show evidence for higher relative survival (SPB unpublished data), roadside wood frogs in these populations lay 10.5% more eggs than woodland populations, countering a portion of the survival disadvantage (Brady 2013). Thus, potential adaptations at adult life history stages and/or investment in offspring quantity over quality might help explain the equivalent population sizes of roadside and woodland demes despite the survival disadvantage of the roadside deme. Relatedly, temporal dynamics of adaptation and maladaptation could also play a role. For example, roadside wood frogs might be maladapted to average road conditions, but adapted to different selection pressures (e.g. disease outbreaks) that vary in time.

Interestingly, within the woodland environment, the roadside deme performed equivalently well as the woodland deme. This indicates that the pattern of local maladaptation is specific to the roadside environment. This departs from previous results showing evidence for deme depression, in which the roadside deme expressed a survival disadvantage in both environments (Brady 2013). Here, that roadside populations have the capacity to survive at higher rates in woodland ponds compared to their natal roadside ponds would suggest that optimally, roadside populations should preferentially breed in woodland ponds, where fitness appears to be relatively higher. That this does not occur suggests that roadside populations might actually be locally adapted at later life history stages, or that roadside environments act as demographic sinks sustained by high rates of poorly conditioned immigrants.

In addition to differential survival, I found an unexpected effect of exposure treatment on both size and development (Fig. 3a, 3c). This effect only occurred in the woodland deme, whereby exposure to spring water increased final development stage and size of larvae grown in the roadside environment. These differences are unrelated to the hypothesis of this study and were inferred from marginally significant effects (*P* = 0.088 and 0.071, respectively) and so should be viewed cautiously. However, these responses suggest that the osmotic environment experienced in the first two days of embryonic development might have carryover effects on growth and development into larval stages for some populations. Further, this initially positive influence of spring water on performance might be reversed at later stages, similar to reports of salt-induced carryover effects on larval growth and juvenile survival in the wood frog (Dananay et al. 2015). Finally, that the roadside deme did not respond in this manner provides further support for differentiation between demes.

Together, these insights into deme level differences should serve as a banner for those studying road effects. Though traditional ecological methods have been invaluable for gaining initial understanding, most of our knowledge of road effects on amphibians has been generated without insights into relative responses of roadside populations (e.g. Karraker et al. 2008; Sanzo and Hecnar 2006). Rather inference into road effects has typically been generated from studies of a single, road-naïve population (e.g. Petranka and Francis 2013), which do not capture how road-affected populations might respond differently owing to evolutionary and plastic effects. These prior studies are of great value. However, because roadside populations are differentiated in their capacity to tolerate road effects, it is critical to infer responses specific to those populations. This is especially poignant here, where inference from woodland populations alone would yield anti-conservative results. Improving our understanding therefore requires we move beyond traditional methods in favor of population specific, evolutionary approaches.

A full understanding of putative local maladaptation in this system will require knowledge of the traits causing the survival disadvantage. As of now, there is no clear evidence of the specific traits that might be influencing this effect. However, given that the only difference detected between the roadside and woodland environment was found to be specific conductance, traits associated with osmotic stress (e.g. gill physiology) and/or contaminant tolerance (e.g. renal function) would be good candidates for future study. Indeed, a body of work highlights a suite of effects induced my osmotic stress (Hua and Pierce 2013; Karraker and Gibbs 2011; Wu et al. 2012) with evidence for physiological mechanisms mediating responses (Wu et al. 2014).

Overall, the putative maladaptation of the roadside deme highlights the value of incorporating evolutionary perspectives into conservation studies by revealing that populations compromised by environmental change can be further compromised by trans-generational effects. Whether this is mediated by evolution or environmental inheritance, the fact remains that the very populations challenged by road effects appear to be the least tolerant of those effects. Further, the message regarding the need to study road effects at the population level should be resounding. Population specific differences are becoming recognized as the rule rather than the exception (Höglund 2009), and the magnitude of difference can be profound. Still, maladaptive outcomes are surprising because organisms often respond adaptively to changing environments (Hereford 2009). Further surprising is that habitat modification can cause maladaptation patterns in some species, but adaptation in others (e.g. Brady 2012; versus Brady 2013). Though underlying mechanisms remain unknown, such opposing outcomes might be mediated by differential scales of gene flow in this system (Richardson 2012), which can affect the response to selection through processes such as migration load (Garcia-Ramos and Kirkpatrick 1997). Regardless of mechanism, these contrasting results highlight the complexity of population level responses to modified environments (Stockwell et al. 2003). Finally, alongside recent reports, putative maladaptation may be an emerging consequence of environmental change that requires careful consideration in conservation (Christie et al. 2012; Darimont et al. 2009; Robertson et al. 2013).

## Acknowledgements

I am grateful to the Leopold Schepp Foundation. I thank D Skelly, S Alonzo, P Turner, M Urban, A Hendry, J Richardson, and R Calsbeek for key guidance and project advice and helpful conversations on the manuscript. I am grateful to S Bolden for extensive support. S Attwood, B and H Bement, E Crespi, E Hall, and J Bushey assisted in the field.

## Compliance with ethical standards

All applicable institutional and/or national guidelines for the care and use of animals were followed.

## Funding

This research was supported by funding from the Mianus River Gorge Preserve Research Assistantship Program, the Hixon Center for Urban Ecology, and National Science Foundation (DEB 1011335).

## Conflict of interest

The author declares that no conflict of interest exists.

